# Using transcranial magnetic stimulation to map the cortical representation of lower-limb muscles

**DOI:** 10.1101/807339

**Authors:** Jennifer L Davies

## Abstract

The aim of this study was to evaluate the extent to which transcranial magnetic stimulation (TMS) can identify discrete cortical representation of lower-limb muscles in healthy individuals. Data were obtained from 16 young healthy adults (12 women, four men; mean [SD] age 23.0 [2.6] years). Motor evoked potentials were recorded from the resting vastus medialis, rectus femoris, vastus lateralis, medial and lateral hamstring, and medial and lateral gastrocnemius muscles on the right side of the body using bipolar surface electrodes. TMS was delivered through a 110-mm double-cone coil at 63 sites over the left hemisphere. Location and size of the cortical representation and the number of discrete peaks were quantified for each muscle. Within the quadriceps muscle group there was a main effect of muscle on anterior-posterior centre of gravity (p = 0.010), but the magnitude of the difference was very small. Within the quadriceps there was a main effect of muscle on medial-lateral hotspot (p = 0.027) and map volume (p = 0.047), but no post-hoc tests were significant. The topography of each lower-limb muscle was complex, displaying multiple peaks that were present across the stimulation grid, and variable across individuals. The results of this study indicate that TMS delivered with a 110-mm double-cone coil could not reliably identify discrete cortical representations of resting lower-limb muscles when responses were measured using bipolar surface electromyography. The characteristics of the cortical representation of lower-limb muscles reported here provide a basis against which to evaluate cortical reorganisation in clinical populations.

---

Transcranial magnetic stimulation (TMS) can be used to study the representation of muscles within the primary motor cortex. Although the extent of somatotopy within the primary motor cortex is debated (Donoghue et al. 1992; Schieber 2001; Schellekens et al. 2018), TMS has revealed alterations in the cortical representation of muscles in several clinical conditions (Liepert et al. 1995; Schwenkreis et al. 2001, 2003; Tsao et al. 2008, 2011b; Schabrun et al. 2009, 2015; Te et al. 2017). This suggests that TMS can identify clinically meaningful differences in cortical representation between groups of individuals.

The majority of work on the representation of muscles within the primary motor cortex has been conducted for muscles of the hand and upper limb. For the lower limb, TMS has been used to quantify the size and the amplitude-weighted centre (centre of gravity [CoG]) of the cortical representation of the resting tibialis anterior muscle (Liepert et al. 1995; Lotze et al. 2003), the resting quadriceps femoris muscle (Schwenkreis et al. 2003), the resting vastus lateralis muscle (Al Sawah et al. 2014), and the active rectus femoris (Ward et al. 2016b, 2016a; Te et al. 2017) and vastii (Te et al. 2017) muscles. No studies have reported the representation of the hamstring or gastrocnemius muscles.

Although TMS has revealed discrete cortical representation of the deep and superficial fascicles of the paraspinal muscles (Tsao et al. 2011a), the extent to which TMS can be used to identify discrete cortical representation of lower-limb muscles is unclear. Only one study has quantified the representation of multiple lower-limb muscles in the same individuals. In statistical analysis, there was no main effect of muscle, suggesting similar cortical representation of the active rectus femoris and vastii muscles (Te et al. 2017). However, the separation between the representation of the three muscles was smaller in individuals with patellofemoral pain than in healthy controls, suggesting that separation between muscles might be a measure of interest (Te et al. 2017). This supports findings in other chronic musculoskeletal pain conditions, where there was reduced distinction in the cortical representation of two back muscles in individuals with low-back pain (Tsao et al. 2011b), and of two forearm extensor muscles in individuals with elbow pain (Schabrun et al. 2015). The extent to which TMS can be used to identify discrete cortical representation of lower-limb muscles therefore warrants further investigation. In addition, recent studies have identified multiple discrete peaks in the cortical representation of a muscle (Schabrun et al. 2015; Te et al. 2017), but no normative data exists on this measure for the representation of lower-limb muscles.

The aim of this study was to evaluate the extent to which TMS can identify discrete cortical representation of lower-limb muscles in healthy individuals. Cortical representation was mapped for seven resting lower-limb muscles involved in control of the knee joint (rectus femoris, vastus lateralis, vastus medialis, medial hamstring, lateral hamstring, medial gastrocnemius, and lateral gastrocnemius) and was quantified using size, CoG, hotspot and number of discrete peaks. These measures were compared between muscles from the same group (quadricep, hamstring, plantar flexor) to evaluate the extent to which TMS can identify discrete cortical representation of lower-limb muscles. These data describe the characteristics of the cortical representation of lower-limb muscles in healthy individuals and provide a basis against which to evaluate reorganisation in clinical populations.

## Methods

This study was carried out at the Cardiff University Brain Research Imaging Centre and was approved by the Cardiff University School of Psychology Ethics Committee.

### Participants

A convenience sample of 18 young healthy adults (13 women, five men; mean [SD] age 23.0 [2.5] years) was recruited from an existing participant database and advertisements placed around Cardiff University. All participants were screened for contraindications to TMS (including history of seizures, neurological injury or head injury) and to ensure that they met the following inclusion criteria: No recent, recurring or chronic pain in any part of the body, no history of surgery in the lower limbs, and not taking any psychiatric or neuroactive medications. The full screening questionnaire is available on the Open Science Framework (https://osf.io/npvwu/). All participants reported that they were right-leg dominant, defined by the leg they would use to kick a ball. All participants attended a single testing session between January and March 2018, and provided written informed consent prior to the start of the experiment. Participants were instructed to have a good night’s sleep the night prior to the experiment, not to consume recreational drugs or more than three units of alcohol on the day of or night prior to the experiment, and not to consume more than two caffeinated drinks in the two hours prior to the experiment.

### Electromyography

Surface electrodes (Kendall 230 series; Covidien, MA) were placed on the following muscles of the right leg according to the SENIAM project guidelines (http://seniam.org/): Rectus femoris, vastus lateralis, vastus medialis, medial hamstring (semitendinosus), lateral hamstring (biceps femoris), medial gastrocnemius, and lateral gastrocnemius. Prior to electrode placement the skin was prepared with exfoliant and alcohol swabs. Data were passed through a HumBug Noise Eliminator (Digitimer, Hertfordshire, UK) and a D440/4 amplifier (Digitimer) where they were amplified x1000 and bandpass filtered (1–2000 Hz; (Groppa et al. 2012)) before being sampled at 6024 samples per second in Signal software (version 6; Cambridge Electronic Designs, Cambridge, UK). Electromyography (EMG) data were stored for offline analysis and viewed in real time using Signal software.

### TMS

TMS was delivered through a 110-mm double-cone coil (Magstim, Whitland, UK) using a single-pulse monophasic stimulator (200^2^, Magstim). Throughout the experiment, a neuronavigation system (Brainsight TMS navigation, Rogue Resolutions, Cardiff, UK) was used to track the position of the coil relative to the participant’s head. The vertex was identified as the intersection of the interaural line and the line connecting the nasion and inion.

### Experimental protocol

Participants were seated and chair height and arm rests were adjusted to optimise comfort. The coil was placed slightly lateral to the vertex, over the left hemisphere. Stimuli were delivered and stimulus intensity was gradually increased until motor evoked potentials (MEPs) were observed in the EMG data. Participants were instructed to stay relaxed, and this was confirmed by visual inspection of the EMG data in real time. For each participant, a stimulus intensity was selected that elicited consistent MEPs in all muscles, but that would be tolerable for the remainder of the experiment. In two participants (one woman, one man) it was not possible to elicit MEPs on resting muscles with a tolerable stimulus intensity, and these participants did not participate further. In the remaining 16 participants, the mean (SD) selected stimulation intensity was 52 (9.3)% maximum stimulator output (range, 38–65% maximum stimulator output).

The neuronavigation software was used to project a 9 x 7 cm grid with 1-cm spacings over the left hemisphere of a representation of the skull that was visible to the experimenter on a monitor. The front-rightmost corner of the grid was positioned 1 cm to the right and 5 cm anterior to the vertex, and the back-leftmost corner was 5 cm to the left and 3 cm posterior to the vertex (Figure 1). This resulted in a total of 63 grid sites. The target grid was centred slightly anterior to the vertex based on previous reports that the CoG of lower-limb muscles is anterior to the vertex (Schwenkreis et al. 2003; Al Sawah et al. 2014; Ward et al. 2016b, 2016a; Te et al. 2017). The target grid was designed to cover a large area to capture the boundaries of the cortical representations. The order of the targets was randomised and each target was stimulated once at the predetermined stimulus intensity. The inter-stimulus interval was at least 5 s. A break of ~5 min was then taken, before this was repeated. The purpose of this break was to avoid long blocks of stimuli during which the participant’s attention or level of arousal might decline. Five sets of stimuli were performed in total. At each target, the experimenter viewed real-time information on the position of the coil and the error from the target position and did not stimulate until there was <2 mm error in coil position. The position of the coil was recorded for each stimulation. The same experimenter (JD) performed TMS for all participants.

**Fig. 1.**
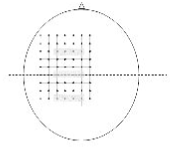
Target stimulation sites. Target stimulation sites were arranged in a 9 x 7 cm grid with 1-cm spacings. The front-rightmost corner of the grid was positioned 1 cm to the right and 5 cm anterior to the vertex, and the back-leftmost corner was 5 cm to the left and 3 cm posterior to the vertex. Grey shading indicates the four stimulation sites (from midline to 3-cm lateral to midline) at the vertex, 3 cm anterior to the vertex, and 3 cm posterior to the vertex over which motor evoked potential latency was averaged for the exploratory analysis

### Data analysis

All processing and analyses were performing using custom-written code in Matlab (versions 2015a and 2019a, MathWorks, Natick, MA, USA). All code used to process and analyse the data is available on the Open Science Framework (https://osf.io/qrsp5/).

#### Planned analyses

Background EMG was quantified as mean absolute EMG in the 250 ms prior to stimulus onset. Trials were automatically discarded if if there was background muscle activity, defined as background EMG greater than three median absolute deviations above the median background EMG from all trials for that muscle. Trials were also excluded if the stimulus was delivered >2 mm from the target. EMG data from the remaining trials were visualised and trials were manually excluded if there were visible artefacts. EMG traces for all muscles of all participants (https://osf.io/3k74p/), and the raw data on which all processing and analyses were performed (https://osf.io/y49uj/) are available on the Open Science Framework (doi 10.17605/osf.io/e7nmk).

EMG data from the remaining trials were averaged across all stimuli at each scalp site. Planned analysis was that the amplitude of the MEP be quantified as the peak-to-peak amplitude of the EMG signal between 10 and 70 ms after stimulus onset. In three participants, this window included the beginning of a second MEP and was, *a posteriori*, shortened to finish at 60 ms after stimulus onset. Visual inspection of EMG data confirmed that the windows captured the MEP for all participants and muscles.

Within each participant and each muscle, MEP amplitude was scaled to peak MEP amplitude (Tsao et al. 2008, 2011b). Map volume was calculated as the sum of scaled MEP amplitudes across all sites (Te et al. 2017). Discrete peaks were identified if the scaled MEP amplitude at a grid site was greater than 50%, was at least 5% greater than the scaled MEP amplitude at all but one of the surrounding grid sites, and was not adjacent to another peak (Schabrun et al. 2015; Te et al. 2017).

The location of each grid site was expressed relative to the vertex. The scaled MEP amplitude, and the medial-lateral (ML) and anterior-posterior (AP) coordinates of the grid site were spline interpolated in two dimensions to obtain a resolution of one millimetre for each axis (Borghetti et al. 2008; Ward et al. 2016a, 2016b). The interpolated data were used to create a topographical map and calculate CoG. CoG is the amplitude-weighted indication of map position, and was calculated using the following formulae:

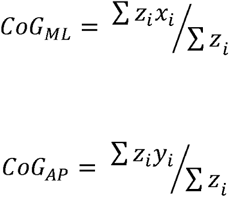

where *x*_*i*_ is the medial-lateral location of the grid site, *y*_*i*_ is the anterior-posterior location of the grid site, and *z*_*i*_ is the scaled amplitude of the MEP at that grid site.

For each outcome and each muscle, the distribution of the data was evaluated using visual inspection of histograms in conjunction with the Anderson-Darling test at the 5% significance level. CoG_ML_, CoG_AP_ and map volume were not different to the normal distribution. Within each muscle group, CoG_ML_, CoG_AP_ and map volume were compared across muscles using a repeated-measures one-way analysis of variance (quadricep muscle group [vastus lateralis, rectus femoris, vastus medialis]) or paired t-test (hamstring [medial, lateral], and plantar flexor [medial gastrocnemius, lateral gastrocnemius] muscle groups). Effect size was calculated as η^2^ for one-way analysis of variance (Tomczak and Tomczak 2014) and *d*_*z*_ for paired t-test (G*Power 3.1 manual, http://www.psychologie.hhu.de/arbeitsgruppen/allgemeine-psychologie-und-arbeitspsychologie/gpower.html; last accessed 15/10/2019). The number of discrete peaks was compared across muscles using a Friedman test (quadriceps) or a Wilcoxon signed-rank test (hamstrings, plantar flexors). The signed-rank test was performed using the approximate method (specific in Matlab software). Effect size was calculated as Kendall’s coefficient of concordance (W) for Friedman test and *r* for signed-rank test (Tomczak and Tomczak 2014).

#### Exploratory analyses

The following analyses were conceived and performed after the data had been viewed.

CoG can be influenced by the presence of multiple discrete peaks in the topography. The stimulation location that elicited the largest MEP (hotspot) was quantified as an additional measure of the cortical representation.

For each participant, the onset latency of the MEP was determined for each stimulation site. Latency was initially defined as the first point after stimulation at which the full-wave rectified EMG signal was more than five median absolute deviations above the median full-wave rectified EMG signal in the 250 ms prior to stimulation and stayed above this threshold for at least 1 ms. The full-wave rectified EMG from each simulation site was then visually inspected to ensure that latency was accurately identified. Latency was averaged across four stimulation sites (from midline to 3-cm lateral to midline) at the vertex (central), 3 cm anterior to the vertex (anterior), and 3 cm posterior to the vertex (posterior; see Figure 1).

In some muscles of some participants, a second MEP was present with a latency of ~60 ms. For each participant and muscle, the presence or absence of a late MEP was determined by visual inspection of the EMG data. The onset latency of this late MEP was determined manually for each muscle by clicking a cursor on a graph where the first deviation from ongoing EMG was visible.

For several muscles, hotspot_ML_ and hotspot_AP_ were different from the normal distribution. Within each muscle group, hotspot_ML_ and hotspot_AP_ were compared across muscles using a Friedman test (quadriceps) or a Wilcoxon signed rank test (hamstrings, plantar flexors). For several muscles, MEP latency at central, posterior, and/or anterior stimulation locations were different from the normal distribution. For each muscle, onset latency was compared across stimulation locations using a Friedman test. Effect sizes were calculated as described above.

## Results

Data were collected from 16 participants (12 women, four men; mean [SD] age 23.0 [2.6] years). All participants completed the testing session and did not report any adverse effects. In six participants, stimulation at the most anterior row of grid sites was uncomfortable due to large twitches of the facial muscles. In one of these participants and one additional participant, stimulation at the most lateral column of grid sites was also uncomfortable due to large twitches in hand muscles. These grid sites were not stimulated for these participants. In the remaining nine participants all 63 grid sites were stimulated.

For one participant (#9), EMG data from the medial gastrocnemius were of poor quality. For another participant (#15), EMG data from the medial gastrocnemius and the lateral hamstring were of poor quality. For a third participant (#7) EMG data from the medial gastrocnemius and medial hamstring were of poor quality. These five muscles (from three participants) were excluded from further analysis.

MEPs were present in all muscles from all participants. CoG and hotspot are shown in Figure 2. MEP latency, CoG, hotspot, map volume and the number of discrete peaks for each muscle are shown in Table 1. Within the quadriceps muscle group there was a significant main effect of muscle on CoG_AP_ (analysis of variance p = 0.010), but the effect size was very small (η^2^ = 0.003). Post-hoc tests showed that CoG_AP_ was more negative (posterior) for vastus medialis than for vastus lateralis (Bonferroni corrected p = 0.035) and rectus femoris (Bonferroni corrected p = 0.018), but the magnitude of this difference was very small (Table 1 and Figure 2A). CoG_AP_ was similar for rectus femoris and vastus lateralis (Bonferroni corrected p > 1).

**Table 1.** Characteristics of the cortical representation of each muscle

|  | Quadriceps |  |  |  |  | Hamstrings |  |  |  | Plantar flexors |  |  |  |
| --- | --- | --- | --- | --- | --- | --- | --- | --- | --- | --- | --- | --- | --- |
|  | Vastus<br>medialis | Rectus<br>femoris | Vastus<br>lateralis | p | Effect<br>size | Medial<br>hamstring | Lateral<br>hamstring | p | Effect<br>size | Medial<br>gastrocnemius | Lateral<br>gastrocnemius | p | Effect<br>size |
| Latency (ms) | 23.9 (1.4) | 21.7 (1.3) | 24 (3.2) | - | - | 25.4 (1.7) | 26.1 (2.2) | - | - | 33.5 (3) | 32.1 (3) | - | - |
| AP CoG (mm) | -2 (9) | -1 (9) | -1 (8) | 0.01 | 0.003 | -1 (9) | -1 (9) | 0.75 | 0.086 | 0 (8) | 1 (8) | 0.30 | 0.086 |
| ML CoG (mm) | -17 (5) | -17 (5) | -17 (4) | 0.33 | 0.000 | -18 (5) | -17 (5) | 0.16 | 0.395 | -17 (5) | -17 (5) | 0.35 | 0.395 |
| AP hotspot (mm)* | -15 (30) | -5 (30) | -10 (25) | 0.18 | 0.106 | -10 (40) | -15 (20) | 0.07 | 0.482 | -10 (32.5) | -10 (40) | 1 | 0.000 |
| ML hotspot (mm)* | -10 (10) | -10 (10) | -10 (5) | 0.03 | 0.225 | -15 (20) | -15 (10) | 0.62 | 0.131 | -10 (22.5) | -20 (12.5) | 0.37 | 0.241 |
| Map volume (%) | 1645 (577) | 1866 (770) | 1789 (656) | 0.05 | 0.014 | 1596 (694) | 1560 (796) | 0.76 | 0.084 | 2085 (960) | 2131 (739) | 0.8 | 0.084 |
| Number of discrete peaks* | 3.5 (2) | 4 (2.5) | 3 (3) | 0.32 | 0.071 | 4 (5) | 3 (3) | 0.96 | 0.012 | 4 (3.3) | 5 (3.3) | 0.07 | 0.481 |

\*Evaluated using non-parametric statistics
Data are mean (standard deviation) for variables evaluated with parametric statistics and median (interquartile range) for variables evaluated with non-parametric statistics. Effect size is $\eta^2$ for one-way analysis of variance, $d_z$ for paired t-test, Kendall's coefficient of concordance (W) for Friedman test and $r$ for signed-rank test.
AP, anterior-posterior; CoG, centre of gravity; ML, medial-lateral.
N = 16 for quadriceps, 14 for hamstrings, and 13 for plantar flexors.

**Fig. 2.**
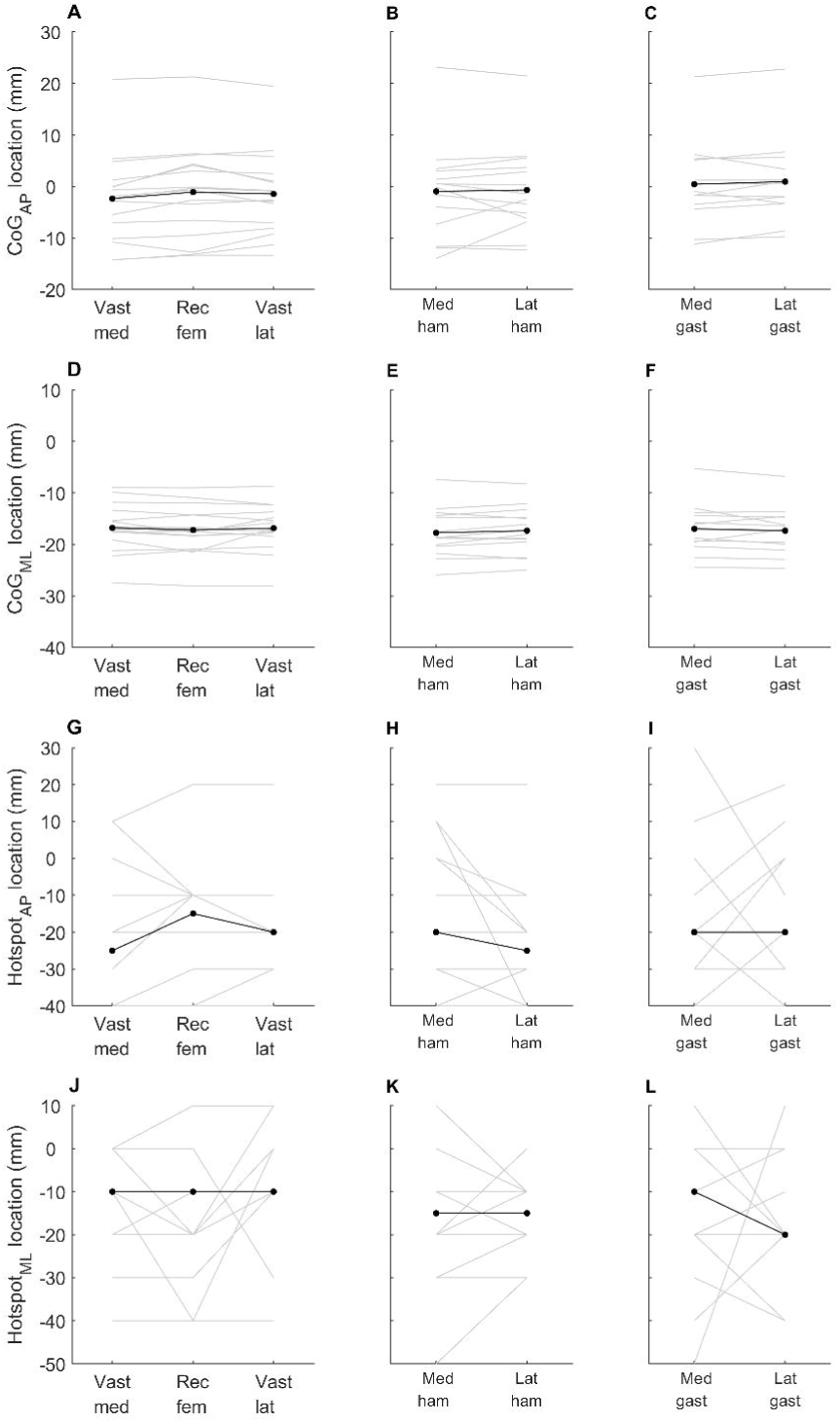
Location of the cortical representation for each muscle. Anterior-posterior (AP) and medial-lateral (ML) centre of gravity (CoG; A–F) and hotspot (G–L) for each muscle in the quadriceps (left column; A, D, G, J), hamstring (centre column; B, E, H, K) and gastrocnemius (right column; C, F, I, L) muscle groups. Grey lines indicate data for individual participants. Black lines indicate mean (for CoG) or median (for hotspot) across all participants. Vast med, vastus medialis; Rec fem, rectus femoris; Vast lat, vastus lateralis; Med ham, medial hamstring; Lat ham, lateral hamstring; Med gast, medial gastrocnemius; Lat gast, lateral gastrocnemius

Within the quadriceps muscle group there was also a significant main effect of muscle on hotspot_ML_ (Friedman test p = 0.027; W = 0.225). No post-hoc tests were significant, and there was no clear trend in the data (Figure 2J). Within the quadriceps muscle group there was a significant main effect of muscle on map volume (analysis of variance p = 0.047, η^2^ = 0.014). No post-hoc tests were significant, but there was a trend for greater map volume in rectus femoris than in vastus medialis (Bonferroni corrected p = 0.07; Table 1). There was no significant effect of muscle for any other outcome measure in any other muscle group (Table 1).

Topographical maps for all muscles from one participant are shown in Figure 3. Topographical maps for the rectus femoris muscle from several participants are shown in Figure 4. These show a complex and variable topography with multiple peaks present across the stimulation grid, including at the boundaries of the grid. Topographical maps for all muscles from all participants are available on the Open Science Framework (https://osf.io/4m2x9/).

**Fig. 3.**
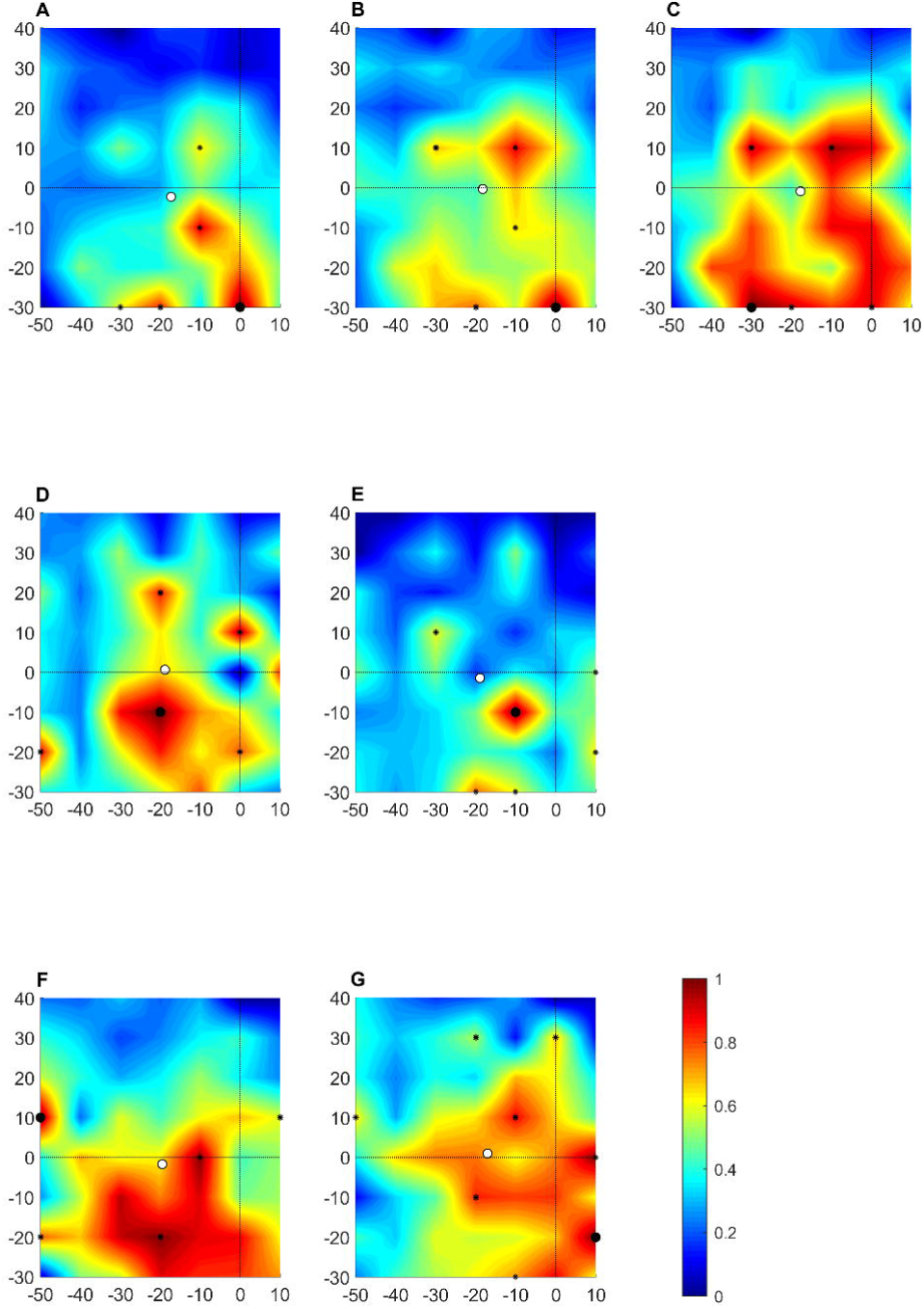
Topographical maps for each muscle from one participant. A: vastus medialis; B: rectus femoris; C: vastus lateralis; D: medial hamstring; E: lateral hamstring; F medial gastrocnemius; G: lateral gastrocnemius. Colour represents scaled amplitude of the motor evoked potential, as indicated in the colour bar. Large black dot indicates the centre of gravity of the cortical representation. Small black dots indicate discrete peaks in the cortical representation. White dot indicates hotspot. Solid black lines indicate the interaural line and the line connecting the nasion and inion

**Fig. 4.**
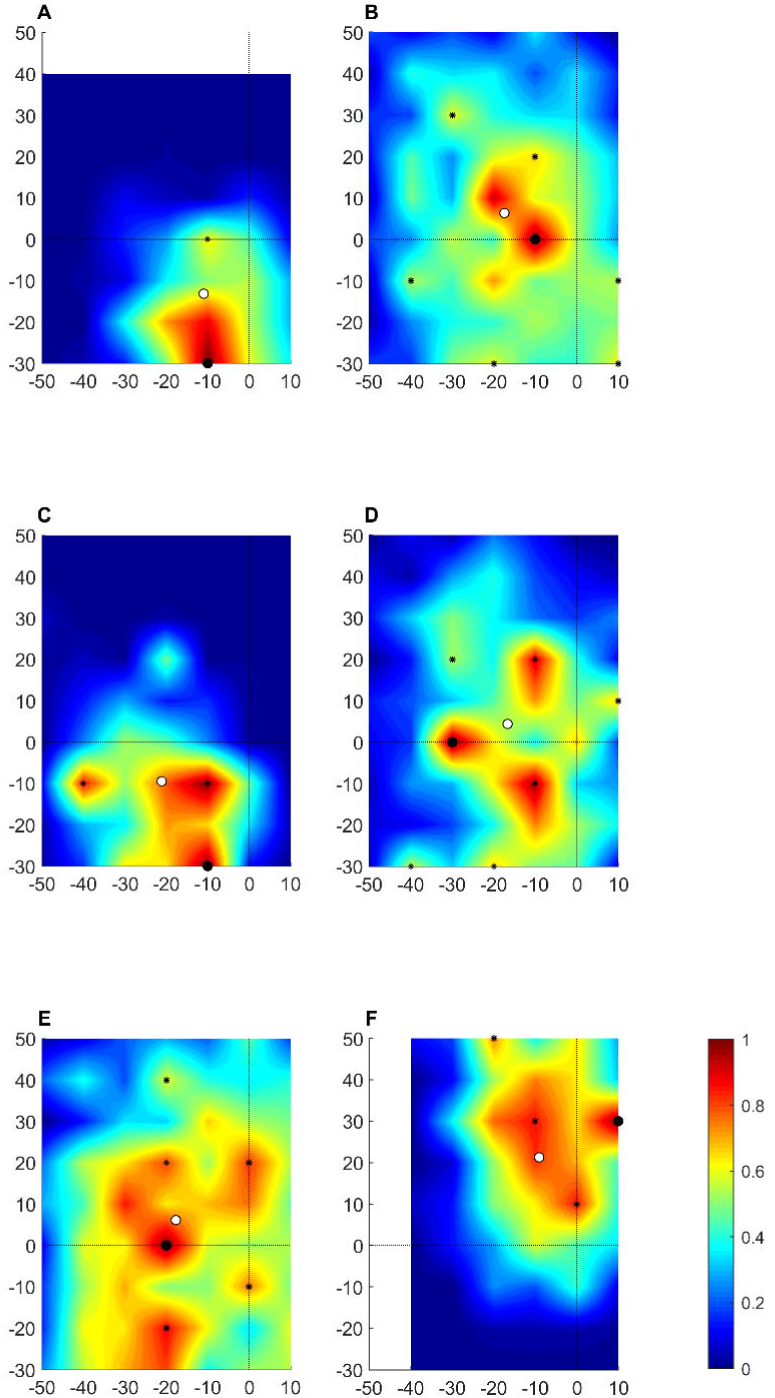
Topographical maps for the rectus femoris muscle from several participants. Large black dot indicates the centre of gravity of the cortical representation. Colour represents scaled amplitude of the motor evoked potential, as indicated in the colour bar. Small black dots indicate discrete peaks in the cortical representation. White dot indicates hotspot. Solid black lines indicate the interaural line and the line connecting the nasion and inion

There was a significant main effect of stimulation location on MEP latency for the vastus medialis (p = 0.03), rectus femoris (p = 0.002), vastus lateralis (p = 0.006) and medial gastrocnemius (p = 0.04; Table 2 and Figure 5). There was no significant main effect for medial hamstring (p = 0.20), lateral hamstring (p = 0.67) or lateral gastrocnemius (p = 0.31; Table 2 and Figure 5). In all muscles, the tendency was for latency to be longer for stimuli delivered at the anterior location than for stimuli delivered at the central and posterior locations. The detailed results of post-hoc tests are provided in Table 2.

**Table 2.** MEP latency at anterior, central and posterior stimulation sites for each muscle

|  | n | Latency (ms) |  |  | p | Effect size (W) | Post-hoc pairwise comparison<br>(corrected p) |  |  |
| --- | --- | --- | --- | --- | --- | --- | --- | --- | --- |
|  |  | Anterior | Central | Posterior |  |  | Anterior-<br>central | Anterior-posterior | Central-<br>posterior |
| Vastus Medialis | 10 | 24.3 (1.6)* | 23.5 (1.5) | 23.5 (1.7) | 0.03 | 0.356 | 0.03 | 0.22 | >1 |
| Rectus Femoris | 12 | 22.4 (2.6)* | 20.7 (2.3)* | 20.8 (1.8) | 0.002 | 0.528 | 0.001 | 0.02 | >1 |
| Vastus Lateralis | 10 | 24.3 (2.7)* | 22.6 (2.5)* | 23.1 (3) | 0.006 | 0.515 | 0.02 | 0.03 | >1 |
| Medial Hamstring | 10 | 25.7 (1.7) | 24.2 (3.4) | 25.7 (4.5) | 0.20 | 0.160 | - | - | - |
| Lateral Hamstring | 10 | 26.3 (2) | 26.0 (1.9) | 25.8 (3) | 0.67 | 0.040 | - | - | - |
| Medial Gastrocnemius | 11 | 36.3 (6.1) | 33.8 (3.3)* | 32.5 (3.5) | 0.04 | 0.298 | 0.25 | 0.06 | 0.16 |
| Lateral Gastrocnemius | 11 | 33.1 (3.5) | 32.1 (2.7) | 32.2 (2.6) | 0.31 | 0.107 | - | - | - |

Latency data are median (interquartile range). Main effect p was obtained from Friedman's test. Post-hoc p were obtained from Wilcoxon signed-rank tests.
Effect size is Kendall's coefficient of concordance.
MEP, motor evoked potential.
\*Different from latency of MEP evoked from stimulation at posterior stimulation sites.

**Fig. 5.**
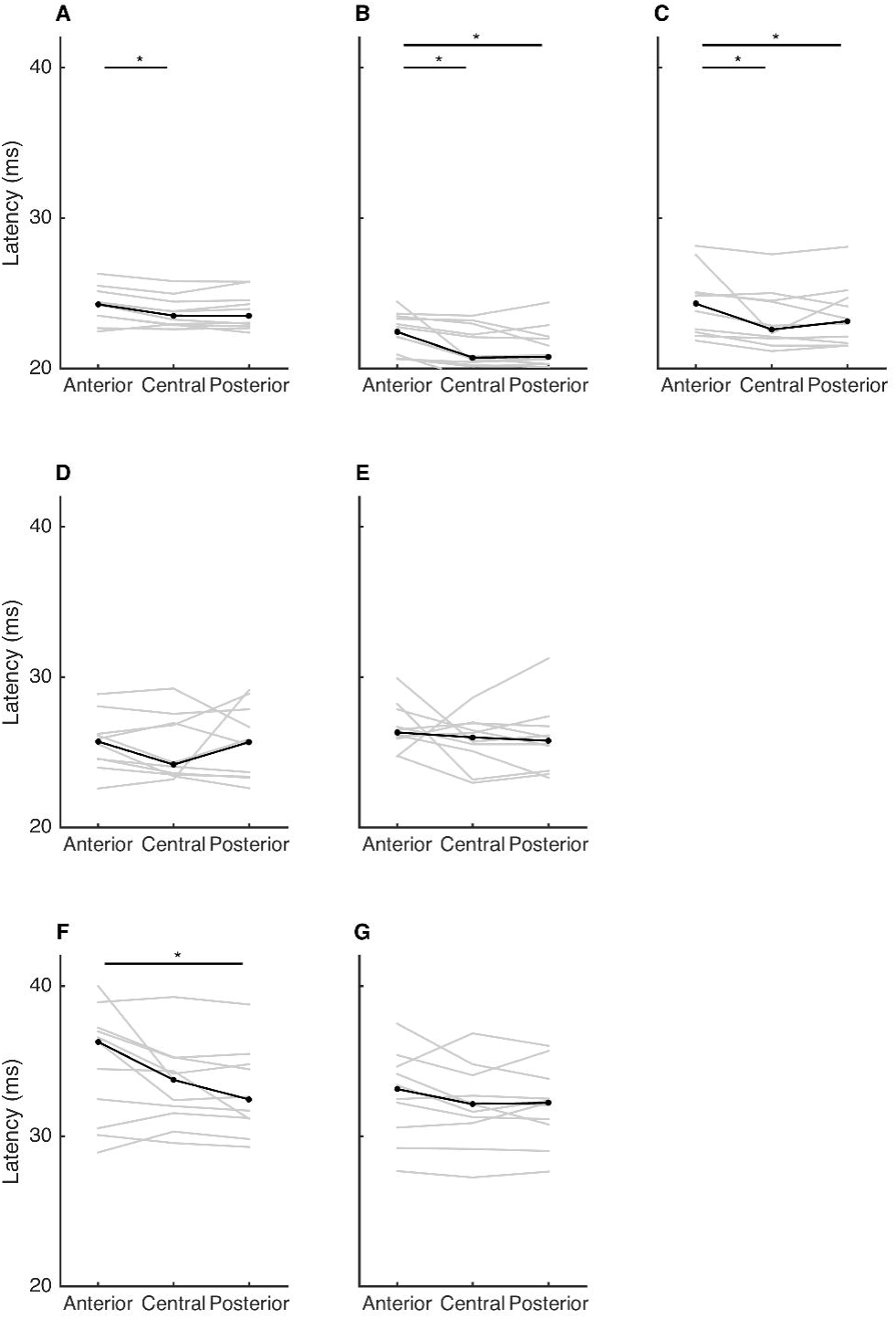
Latency of motor evoked potential evoked from stimulation at anterior, central and posterior stimulation sites. Data are shown for the quadriceps (top row), hamstring (middle row) and gastrocnemius (bottom row) muscles. A: vastus medialis; B: rectus femoris; C: vastus lateralis; D: medial hamstring; E: lateral hamstring; F medial gastrocnemius; G: lateral gastrocnemius. Grey lines indicate data for individual participants. Black lines indicate median across all participants. Asterisks indicate significant difference (p < 0.05).

EMG data from the medial and lateral hamstring muscles from one participant are shown in Figure 6. In these muscles, a second MEP was present after the primary MEP. This late MEP was present in the lateral hamstring of five participants, the medial hamstring of four participants, the vastus medialis and rectus femoris of two participants and the vastus lateralis of one participant. The late MEPs observed in the quadriceps muscles were all very small, whereas those observed in the hamstring muscles could be sizeable (see Figure 6). The mean (SD) estimated onset latency for the late MEP was 61 (3) ms for the lateral hamstring (n = 5), 62 (7) ms for the medial hamstring (n = 4), 68 (2) ms for the vastus medialis (n = 2), 67 (0) ms for the rectus femoris (n = 2) and 65 ms for the vastus lateralis (n = 1).

**Fig. 6.**
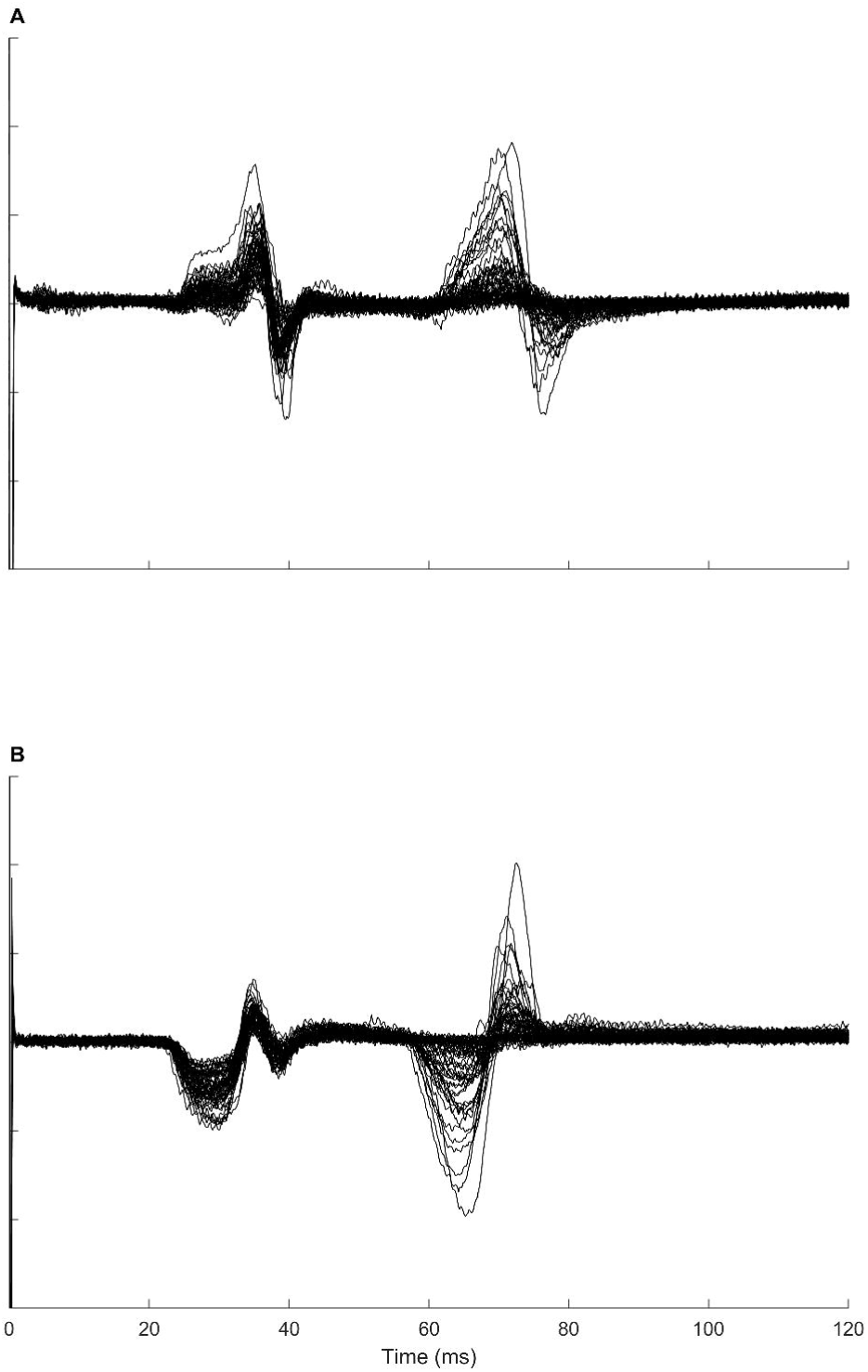
Late motor evoked potential. Surface electromyography data from the medial (A) and lateral (B) hamstring of one participant. A late motor evoked potential is clearly evident with a latency of approximately 60 ms.

## Discussion

In this study I used TMS to map the cortical representation of seven resting lower-limb muscles in healthy individuals. The data indicate that the size, CoG, hotspot and number of discrete peaks were largely similar across muscles within each group (quadriceps, hamstrings, plantar flexors). There was a statistically significant different in CoG_AP_ and hotspot_ML_ across the quadriceps muscles but the effect size and magnitude of differences was very small. The magnitude of the difference means that it would not be practically possible to differentially target one of the three quadriceps muscles with navigated TMS. These results indicate that there was considerable overlap in the cortical representations of the lower limb muscles identified by TMS and surface EMG, and that TMS delivered with a double-cone coil could not identify discrete cortical representations of lower-limb muscles when MEPs were measured with bipolar surface EMG.

The topography of the seven lower-limb muscle studied here was often complex, displaying multiple peaks that were present across the stimulation grid, and variable across individuals. This may reflect a large and complex anatomical representation of these muscles within the cortex, with considerable inter-individual variability. However, the impact of the techniques used to quantify the cortical representation must also be considered, particularly the volume of cortical tissue excited by the TMS and the potential for peripheral volume conduction (crosstalk) in the surface EMG recordings.

For all muscles, the average CoG was located at, or slightly posterior to, the vertex. In addition, despite the large area covered by the target grid, large MEPs were often observed in response to stimuli delivered at the edge of the grid. This is in contrast to previous mapping studies of the quadriceps muscles, which have reported an anterior CoG (Schwenkreis et al. 2003; Al Sawah et al. 2014; Ward et al. 2016b, 2016a; Te et al. 2017) and relatively constrained map boundaries (see Figure 1 in (Te et al. 2017) and Figure 2 in (Ward et al. 2016a)). These previous studies have all used a figure-of-eight stimulation coil, in contrast to the double-cone coil used here, and some (Ward et al. 2016b, 2016a; Te et al. 2017) have studied active muscles, in contrast to resting muscles studied here.

### Double-cone vs. figure-of-eight coil

The double-cone coil used in this experiment consists of two circular coils arranged in a figure-of-eight shape with a fixed angle of about 95 degrees between the two wings. The double-cone coil can stimulate deeper regions of the brain than a circular or figure-of-eight coil, which is advantageous for activating corticospinal projections to the lower-limb muscles. It elicits MEPs at a lower stimulation intensity than a circular coil and a figure-of-eight coil (Dharmadasa et al. 2019). However, this improved stimulation depth is obtained at the expense of focality (Lu and Ueno 2017). The large area of cortical tissue activated by TMS delivered through a double-cone coil means that even when the coil is not centred over the motor cortex, excited tissue will likely include the motor cortex. This could explain the apparent expansion of the cortical representation beyond the borders of the target grid, and the larger area of the cortical representation than previously reported (Schwenkreis et al. 2003; Al Sawah et al. 2014; Ward et al. 2016b, 2016a; Te et al. 2017). Nonetheless, if current spread was the only relevant factor, the largest MEP may still be expected when the stimulating coil was centred over the motor cortex. The discrete peaks in the topographical maps that occurred at or close to the boundaries of the target grid in some participants suggest a complexity that is difficult to describe solely by current spread.

It is possible that the double-cone coil excited cortical tissue beyond the motor cortex, and this resulted in MEPs. The latency of MEPs observed in response to stimulation at the anterior of the target grid was often slightly longer than that of MEPs observed in response to stimulation at the centre or posterior of the target grid. This may suggest that a different pathway was involved in the generation of these MEPs. Corticospinal neurones innervating the lower limb spinal motoneurones are present in the premotor cortex (He et al. 1993) and supplementary motor area, caudal cingulate motor area on the dorsal bank and the rostral cingulate motor area (He et al. 1995), as well as the primary motor cortex. However, it is also possible that the longer latency at anterior stimulation sites is an artefact of a smaller MEP size at the grid boundary, and this requires further investigation.

The figure-of-eight coil provides a very focal stimulation in superficial cortical regions, but minimal stimulation at increasing depths (Lu and Ueno 2017). If much of the cortical representation of lower-limb muscles is inaccessible to the figure-of-eight coil, the resulting topography will appear less complex than that obtained with a double-cone coil. Although the double-cone coil will excite deeper cortical tissue, the larger current spread means that it also excites a large volume of cortical tissue at shallow depths. It is possible that the double-cone coil accessed corticospinal neurones that could not be accessed with the figure-of-eight coil, and thus the complexity of the cortical representation reflects the true anatomical complexity that cannot be uncovered with a figure-of-eight coil.

### Surface EMG

High-density surface EMG recordings suggest that MEPs recorded in forearm muscles using conventional surface EMG may contain crosstalk from neighbouring muscles (van Elswijk et al. 2008; Gallina et al. 2017; Neva et al. 2017). No analogous studies have been performed in lower-limb muscles. The identification of crosstalk in voluntary contractions without using high-density surface EMG is difficult, as recently highlighted by Talib et al. (Talib et al. 2019). Studies with high-density surface EMG are required to evaluate the influence of crosstalk on MEPs evoked from lower-limb muscles. Bipolar surface EMG has been used to map the representation of lower-limb muscles in healthy (Weiss et al. 2013; Ward et al. 2016a) (Liepert et al. 1995; Lotze et al. 2003; Schwenkreis et al. 2003; Al Sawah et al. 2014; Ward et al. 2016a; Te et al. 2017) and clinical (Ward et al. 2016b; Te et al. 2017) populations. The current results indicate bipolar surface EMG used with TMS delivered through a double-cone coil cannot reliably identify discrete cortical representation of lower-limb muscles in young, healthy individuals.

### Resting vs. active muscle

If, as has been suggested, the cortical representation of muscles is functional, rather than anatomical (Schieber 2001; Ejaz et al. 2015; Leo et al. 2016), then requiring the participant to perform a motor task will engage the specific subset of cortical neurones relevant for that task. The cortical representation revealed by TMS may then be biased towards the representation for that specific task, at the expense of the representations for other functions of the target muscle. For example, requiring the participant to perform an isometric contraction of the quadriceps would increase excitability of the cortical areas involved in generating this type of contraction. The cortical representation of the quadriceps revealed by TMS will reflect this, and may fail to include the cortical representation relevant for a dynamic movement such as gait. For this reason, I chose to study resting muscles in the present study. Ward et al. (Ward et al. 2016b, 2016a) and Te et al. (Te et al. 2017) studied muscles performing a low-intensity isometric contraction, and this may have contributed to the more focussed topographical maps in these previous studies.

By contrast, Schwenkreis et al. (Schwenkreis et al. 2003) and Al Sawah et al. (Al Sawah et al. 2014) studied resting muscles. The topographical maps are not described in detail, but Schwenkreis et al. report that, across six healthy subjects, they observed an MEP in the resting quadriceps muscle response to stimulation at, on average, ~15 sites (*cf* map area results) (Schwenkreis et al. 2003). Similarly, Al Sawah observed an MEP in the resting vastus lateralis muscle in response to stimulation at, on average, ~8 sites (n = 10 healthy participants, *cf* map area results) (Al Sawah et al. 2014). The size of these MEPs, and whether the topography incorporated multiple discrete peaks, is not clear. However, the available data suggest that the cortical representations uncovered were smaller and less complex than those revealed in the present study. This suggests that the difference between the current and previous studies cannot be explained by the state of the muscle, and is more likely a function of the stimulating coil.

### Late MEPs

In some muscles in some participants, there was a second MEP that occurred with a latency of 60–70 ms. This late MEP has previously been reported in the resting hamstrings, quadriceps, tibialis anterior and triceps surae muscles, with a latency of 59 ms, 64 ms, 79 ms and 72 ms, respectively (Dimitrijević et al. 1992), and in resting and active tibialis anterior and triceps surae muscles with a latency of ~100 ms (Holmgren et al. 1990). Dimitrijevic et al. reported that this late MEP was most prevalent in the hamstrings, where it was of higher amplitude than the primary MEP. This is in line with the current findings, where the late MEP was observed most frequently and with the largest amplitude in the hamstring muscles. The late MEP is not exclusive to the lower limbs, and has been observed in resting and active forearm muscles (Holmgren et al. 1990) and a resting, but not active, intrinsic hand muscle (Wilson et al. 1995).

The source of the late MEP is not known, and could be central or peripheral. Indirect cortico-spinal or cortico-brainstem-spinal pathways, which originate either from the targeted areas of the motor cortex or from wider cortical areas excited by the stimulation, could play a role. Proprioceptive information arising from the primary MEP could also play a role. Based on the latency of responses from several lower-limb muscles, Dimitrijevic et al. argued against a segmental or transcortical stretch reflex origin of the late MEP, and against the involvement of gamma motor neurones (Dimitrijević et al. 1992). The difference in the latency of the late MEP between upper- and lower-limb muscles is ~5 ms greater than the difference in latency of the early, primary MEP (Holmgren et al. 1990), lending the possibility that a slow central pathway is involved. Recent evidence indicates the primary motor cortex includes slow pyramidal tract neurones (Innocenti et al. 2019), which comprise the majority of the pyramidal tract but are not well studied (Firmin et al. 2014; Kraskov et al. 2018). This may provide one such candidate pathway, but this remains to be studied.

### Limitations

Responses to TMS were evaluated using bipolar surface EMG, and the results may have been influenced by peripheral volume conduction (crosstalk) from other muscles. Studies using high-density surface EMG recordings, similar to those conducted in the forearm muscles (van Elswijk et al. 2008; Gallina et al. 2017; Neva et al. 2017), are required to elucidate the contribution of cross-talk to MEPs recorded from lower-limb muscles. However, the finding that TMS could not identify discrete cortical representations of lower-limb muscles measured with bipolar surface EMG is highly relevant, and all TMS studies of the lower-limb to date have been performed using bipolar surface EMG.

TMS was delivered through a double-cone coil. The ability of other coil designs, such as the figure-of-eight coil, to identify discrete cortical representations of lower-limb muscles remains to be determined. When using any coil, the volume of cortical tissue excited by the stimulus, and whether this is likely to encompass the full cortical representation of the target muscle, should be considered. This is particularly relevant for flat coils, where the depth of electric field penetration is lower than for the double-cone coil (Lu and Ueno 2017). The current results are particularly relevant in light of a recent study recommending the standard use of the double-cone coil for lower-limb studies, in preference to a figure-of-eight or circular coil (Dharmadasa et al. 2019).

### Conclusions

The results of this study indicate that TMS delivered with a 110-mm double-cone coil cannot reliably identify discrete cortical representations of resting lower-limb muscles when MEPs are measured using bipolar surface EMG. The characteristics of the cortical representation of lower-limb muscles reported here provide a basis against which to evaluate cortical reorganisation in clinical populations.

## Acknowledgements

J Davies is funded by the Biomechanics and Bioengineering Research Centre Versus Arthritis at Cardiff University. Funding for this project was provided by a Wellcome Trust Institutional Strategic Support award from Cardiff University to J Davies.

